# NAD consumption by PARP1 in response to DNA damage triggers metabolic shift critical for damaged cell survival

**DOI:** 10.1101/375212

**Authors:** Michael M. Murata, Xiangduo Kong, Emmanuel Moncada, Yumay Chen, Ping Wang, Michael W. Berns, Kyoko Yokomori, Michelle A. Digman

## Abstract

DNA damage signaling is critical for the maintenance of genome integrity and cell fate decision. Poly(ADP-ribose) polymerase 1 (PARP1) is a DNA damage sensor rapidly activated in a damage dose and complexity-dependent manner playing a critical role in the initial chromatin organization and DNA repair pathway choice at damage sites. However, the cell-wide consequence of its activation in damaged cells is not well delineated. Using the phasor approach to fluorescence lifetime imaging microscopy (FLIM) and fluorescence-based biosensors in combination with laser microirradiation, we found a rapid cell-wide increase of the bound/free NADH ratio in response to nuclear DNA damage, which is triggered by NAD+ depletion by PARP activation. This change is linked to the metabolic balance shift to oxidative phosphorylation (oxphos) over glycolysis. Inhibition of the respiratory chain resulted in rapid PARP-dependent ATP reduction and intracellular acidification, and eventually, PARP-dependent, AIF-independent, apoptosis indicating that oxphos becomes critical for damaged cell survival. The results reveal the novel pro-survival effect of PARP activation through a change in cellular metabolism, and demonstrate how unique applications of advanced fluorescence imaging technologies in combination with laser microirradiation can be a powerful tool to interrogate damage-induced metabolic changes at high spatiotemporal resolution in a live cell.

## Introduction

Poly(ADP-ribose) polymerase 1 (PARP1) functions as a DNA damage sensor whose enzymatic activity is rapidly activated in response to DNA damage (1). PARP1 uses oxidized nicotinamide adenine dinucleotide (NAD+) as a substrate to ADP(ribosyl)ate itself and various target proteins. Although initially recognized as a base excision repair (BER) factor, it is now known that PARP1 is involved in multiple DNA repair pathways. Rapid poly(ADP-ribose) (PAR) accumulation at damage sites controls damage site accessibility through the recruitment of various chromatin modifiers and dictates DNA repair pathway choice (1–15). For example, we demonstrated that PARylation at damage sites suppresses the 53BP1 recruitment to damage sites, providing another explanation for 53BP1- and NHEJ-dependent cytotoxicity by PARP inhibition (16).

In addition to promoting efficient repair and thus cell survival, PARP was also shown to affect metabolism and mediate senescence and cell death (both apoptosis and necrosis) when hyperactivated (1). Since the oxidized and reduced forms of NAD (NAD+ and NADH, respectively) are metabolic cofactors critical for cellular energy production (17), consumption of NAD+ by damage-induced PARP activation was expected to hinder cellular energy metabolism. In addition, PARP1 inhibits glycolysis through PAR binding to hexokinase (HK), a critical enzyme in the pathway (18–20), resulting in ATP deprivation and subsequent cell death termed parthanatos (21). Parthanatos also requires PAR-dependent nuclear translocation of apoptosis-inducing factor (AIF) from mitochondria (22). PARP activation was also shown to cause intracellular acidification, which appears to promote necrosis (23). Thus, while the downstream effects of PARP activation may be complex, collectively, it was suggested to trigger cell death as the default result of PARP “hyperactivation”. However, the precise relationship between the DNA damage dosage and the strength of PARP signaling that affects energy metabolism and/or triggers cell death is unclear, and whether or not damage-induced PARP activation has any cell-wide pro-survival effects distinct from its immediate role at DNA damage sites has not been determined.

Because NADH is an essential cofactor for oxidation-reduction (redox) reactions and production of ATP, multiphoton microscopy techniques to capture NADH autofluorescence and measure metabolic dynamics in living cells have recently garnered attention for their broad applicability in studies ranging from development to cancer and aging (24–26). The free and protein-bound states of NADH were shown to reflect glycolysis and oxidative phosphorylation (oxphos), respectively, which can be differentiated using fluorescence lifetime imaging microscopy (FLIM). One of the major challenges of FLIM is the fitting routine required to dissect the possible lifetime contributions of different fluorescent species in a single pixel. In our approach, we apply the fit-free phasor approach to FLIM analysis that provides a graphical representation of the fluorescence decay at each pixel (27). Pixels that have multiple contributions of fluorophore lifetimes can be resolved as a linear combination of the individual species. Furthermore, the absolute concentration of NADH can be measured using the phasor-FLIM method (28), which is expected to sensitively respond to the changes of NAD+ (29). Here, using the phasor-FLIM approach in conjunction with fluorescent biosensors and laser microirradiation, we examined the real time metabolic changes in response to DNA damage with high spatiotemporal resolution and interrogated the contribution of differential PARP signaling in response to different dosage/complexity of DNA damage. Using this comprehensive approach, we observed rapid reduction of NADH (correlating with the decrease of NAD+) and ATP in both the nucleus and cytoplasm in a PARP1 activity-dependent manner. Interestingly, PARP1 activation also triggers a rapid increase of protein-bound NADH species over free NADH, which correlates with the net increase of oxphos over glycolysis. Importantly, this shift appears to reflect the increased metabolic dependence of damaged cells on oxphos. Inhibitors of oxphos highly sensitized cells to DNA damage with exacerbated ATP deprivation, resulting in AIF-independent apoptosis. This increased dependence on oxphos is rescued by PARP inhibition or NAD+ replenishment, and thus, is distinct from PARP-dependent HK inhibition in glycolysis. Taken together, the phasor approach to FLIM and fluorescence biosensors in combination with laser microirradiation provides valuable tools to capture high-resolution single cell dynamics of metabolic changes in response to DNA damage and uncovered the key downstream consequence of NAD+ consumption by PARP1 that promotes cell survival in DNA damage cells.

## Materials and Methods

### Cell culture

HeLa adenocarcinoma cells were cultured in DMEM (Invitrogen, Carlsbad, CA) supplemented with 10% fetal bovine serum (FBS), 2 mM L-glutamate, and 1% penicillin-streptomycin. HCT116 colorectal carcinoma cells were cultured in ATCC-formulated McCoy’s 5a Medium Modified (ATCC 30-2007) supplemented with 10% FBS and 1% penicillin-streptomycin. HFF-1 human foreskin fibroblast cells were cultured in DMEM (ATCC 30-2002) supplemented with 15% FBS and 1% penicillin-streptomycin.

### Inhibitor treatment

Cells were treated for 1 hr before experiments with PARP inhibitor (PARPi) (100 µM NU1025 (Sigma) or 20 µM olaparib (Sigma)), or ATM inhibitor (10 µM KU55933 (Calbiochem)) and DNA-PK inhibitor (10 µM NU7026 (Sigma)), in 0.1% DMSO (Sigma). For respiration inhibition, cells were treated for 1 hr before experiments with the mitochondrial complex I inhibitor rotenone (Sigma, R8875) (1 µM for laser damage and 0.25 µM for MMS or H_2_O_2_-induced damage) and complex III inhibitor antimycin A (Sigma, A8672) (1 µM for laser damage and 0.25 µM for MMS or H_2_O_2_-induced damage) in 0.2% DMSO. For nicotinamide (NAM) treatment, cells were treated 1 hr before laser-induced DNA damage induction with 1 mM NAM (Sigma, 72340). For NAMPT inhibition, cells were treated with 10 nM FK866 (Sigma, F8557) for 6 hr before experiments.

### MMS and H_2_O_2_ treatment

For comparison to the effects of laser-induced DNA damage, cells were treated with 1 mM or 3 mM methyl methanosulfonate (MMS) for 1 hr or 500 µM hydrogen peroxide (H_2_O_2_) for 30 min prior to metabolic and cytotoxic analyses.

### Immunofluorescent staining

Immunofluorescent staining (IFA) was performed essentially as described previously (34). Antibodies used are mouse monoclonal antibodies specific for γH2AX (05–636, Millipore), PAR polymers (BML-SA216–0100, Enzo Life Sciences, Inc.), Mre11 (GTX70212, Gene Tex, Inc.), Actin (Sigma) or cyclobutane pyrimidine dimer (CPD) (MC-062, Kamiya Biomedical Company) as well as rabbit polyclonal antibodies specific for AIF antibody (GTX113306, Gene Tex, Inc.). Affinity-purified rabbit anti-PARP1 antibody was previously described (16,30).

### Confocal fluorescence microscope

All experiments were performed on an inverted confocal Zeiss LSM710 (Carl Zeiss, Jena, Germany). A 40x 1.2NA water-immersion objective (Zeiss, Korr C-Apochromat) was used.

### NIR laser microirradiation

A pulsed titanium-sapphire 100 femtosecond laser (Spectra-Physics, Santa Clara, CA) at 80 MHz tuned for 780 nm two-photon microirradiation was used at the input power of 17.7 mW, 19.9 mW, and 24.6 mW measured after the objective for low, medium, and high damage conditions, respectively. Image sizes of 512×512 pixels were obtained with defined regions of 45×5 pixels for microirradiation. Microirradiated regions were scanned once with a scan speed of 12.61 µs per pixel for damage induction.

### Phasor approach to FLIM and NADH intensity/concentration measurement

Following DNA damage induction, either by laser microirradiation or chemical agents (MMS or H_2_O_2_), 40 frames of FLIM data were acquired with 740 nm two-photon excitation at approximately 2 mW by an ISS A320 FastFLIM box (ISS, Champaign, IL) with image sizes of 256×256 pixels and a scan speed of 25.21 µs per pixel. Fluorescence excitation signal was separated with a dichroic filter (690 nm) and fluorescence was detected by photomultiplier tubes (H7422P-40, Hamamatsu, Japan) with a blue emission filter (495LP) of 420-500 nm to capture NADH auto-fluorescence. FLIM data was transformed into pixels on the phasor plot using the SimFCS software developed at the Laboratory for Fluorescence Dynamics, University of California, Irvine as described previously (27). Coumarin-6, which has a known single exponential lifetime of 2.5 ns, was used as a reference for the instrument response time. Nuclear and cytoplasmic compartments were analyzed separately based on the intensity of NADH signal as previously reported (31). The concentration of NADH was calculated by calibrating with a known concentration of free NADH and correcting for the difference in quantum yield between the free and bound forms of NADH as described previously (28).

### NAD+ measurement

The HeLa cell lines stably expressing either the cytoplasm-or mitochondria-targeted NAD+ sensor, or the corresponding cpVenus controls were kindly provided by Dr. Xiaolu Cambronne (the Oregon Health and Science University) (32). Cells were plated on a glass-bottomed imaging dish and were incubated at 37°C for ~40 hr until they reached 60-80 % confluency. Live images were captured before and 1 hr after indicated treatment. The fluorescence ratios (488 nm/405 nm) were measured and analyzed as previously described (32).

### ATP measurement

HeLa cells were transfected using Lipofectamine 3000 (Invitrogen) with either AT1.03 (cytoplasm-localized) or Nuc AT1.03 (nucleus-localized) plasmids (33). After 18-24 hr, cells were subjected to laser microirradiation or MMS treatment in the presence of DMSO or indicated inhibitors, and the YFP/CFP fluorescence ratios were measured as previously described (34). Experiments are replicated at least three times and 20-30 cells were examined in total.

### Intracellular pH (pHi) measurement

Changes in intracellular pH were measured using the commercial pHrodo Green AM Intracellular pH Indicator kit (ThermoFisher, P35373) according to manufacturer’s instructions.

### Measurement of Mitochondria Respiration: the Seahorse assay

HeLa cells were plated in a 24-well Seahorse XF-24 assay plate at 1 × 10^5^ cells/well and grown for approximately 20 hr. Cells were then incubated with 3 mM MMS for 1 hr in the presence of DMSO or PARP inhibitor. At 1 hr after the release from MMS treatment, Seahorse analysis was performed using a Agilent Seahorse XF24 Extracellular Flux Analyzer following the manufacturer’s protocol as described previously (35).

### Detection of senescence, apoptosis and necrosis and assessment of cytotoxicity

Cellular senescence was determined by β-gal staining as described previously (36). Apoptosis and necrosis was detected using the commercial kit (AB176749, Abcam, Cambridge, England) following manufacturer’s instruction. Cytotoxicity was also examined by propidium iodide permeability assay. After damage induction and inhibitors treatment, propidium iodide and Hoechst 33342 were added into the medium to achieve final concentrations of 0.8 µg/mL and 0.5 µg/mL, respectively. The cells were then incubated at 37 °C for 5-15 min and visualized for the total cells (blue) and dead cells (red) under a confocal fluorescence microscope.

## Results

### Phasor-approach to FLIM reveals ratiometric increase of protein-bound NADH over free NADH in response to DNA damage

Using laser microirradiation, DNA damage-induced changes of the free and bound fractions of NADH were measured by FLIM in the cytoplasm and nucleus in HeLa cells. Laser microirradiation can be used to induce DNA damage in a highly controlled fashion (16,37–40). In particular, titration of the input power of near-infrared (NIR) laser allows differential DNA damage induction, from relatively simple strand breaks to complex DNA damage (i.e. single and double strand breaks (SSBs and DSBs), thymine dimer and base damage) with differential H2AX phosphorylation (γH2AX) and PARP activation (Supple Fig. S1A; Fig. 2A) (16,39). With high input-power, over 70% of irradiated cells were viable after 24 hr and surviving cells were arrested in interphase with no evidence for senescence at least up to 60 hr post irradiation (Supple Fig. S1B and C). Over 90% of cells were viable at 24 hr after microirradiation with medium input power whereas cells actually recovered and proliferated with low input-power irradiation (Supple Fig. S1B).

**Figure 2.**
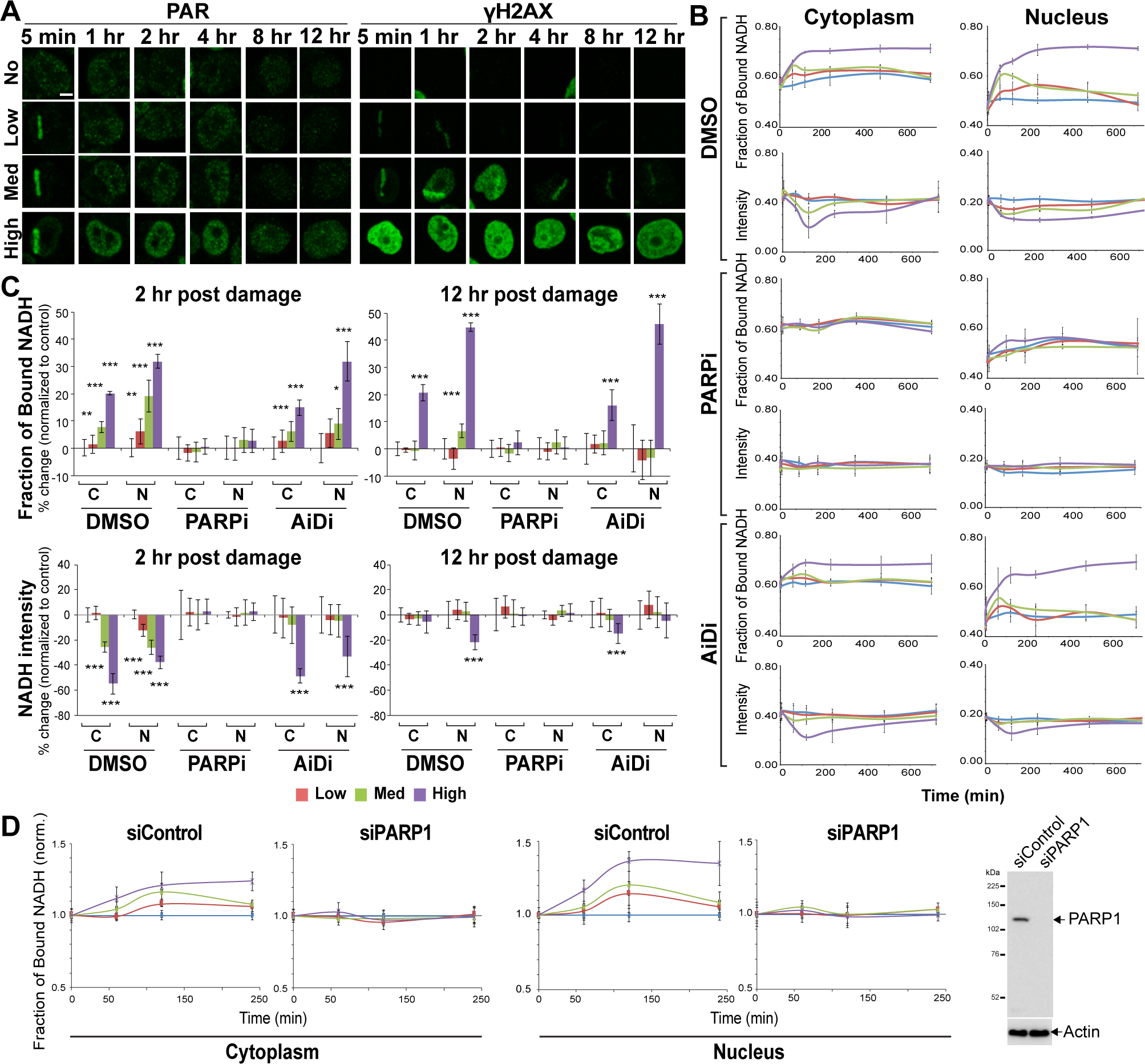
Damage-induced increase of the bound to free NADH ratio is PARP1-dependent. (A) Immunofluorescent staining for PAR (left) and γH2AX (right) in HeLa cells fixed at various time points up to 12 hr post damage following low, medium, and high input laser power. Scale bar = 5 µm. (B) The fraction of bound NADH and intensity of NADH over time in the cytoplasmic and nuclear compartments of HeLa cells treated with either 0.1% DMSO (top), 20 µM PARP inhibitor (olaparib) (middle), or 10 µM ATM inhibitor (KU55933) and 10 µM DNA-PK inhibitor (NU7026) (bottom). (C) The percent change in the fraction of bound NADH (top) and the intensity of NADH (bottom) at 2 hr (left) and 12 hr (right) post damage relative to basal conditions. * p < 0.05, ** p < 0.01, *** p < 0.001. (D) The fraction of bound NADH over time in the cytoplasmic (left) and nuclear (middle) compartments of HeLa cells with siPARP1 or control siRNA. Western blot analysis (right) of siControl and siPARP1 transfected HeLa cells. The whole cell extracts were run on SDS-PAGE and blotted with anti-PARP1 antibody. Anti-actin antibody served as loading control.

Using these varying laser input powers, we examined the effect of nuclear DNA damage on cellular metabolism in real time. Clusters of pixels were detected on the phasor plot and used to pseudo-color the intensity images according to fluorescence lifetime (Fig. 1A and B). We also measured both NADH intensity and concentration (28), which are largely concordant with each other and are reduced in a damage dose-dependent manner (Supplemental Fig. S2A). This is in agreement with the reduction of NAD+ in the mitochondria and cytoplasm as detected by the NAD+-specific biosensor (32) (Supplemental Fig. S2C). FLIM analyses revealed the increase in the ratio of the bound NADH fraction over free NADH within the first 120 min post damage induction in both the cytoplasm and nucleus (Fig. 1C and E). Interestingly, while this increase was transient for low and medium damage conditions, the fraction of bound NADH remained high for over 12 hr with high input power damage, which is particularly prominent in the nucleus, despite the recovery of NADH intensity/concentration (Figs. 1C and 2B; Supplemental Fig. S2A). The results raise the possibility that the shift to bound NADH is not the mere reflection of NADH depletion but represents more long-term metabolic reprogramming in response to damage. Treatment with alkylating (MMS) and oxidizing (H_2_O_2_) damaging agents exhibited similar transient shifts toward a high fraction of bound NADH (Fig. 1D and E). Comparable kinetics of NADH intensity/concentration reduction consistent with NAD+ reduction was also observed (Supplemental Fig. S2B and C). Taken together, the results demonstrate that the increase of bound NADH is a general response to DNA damage, not restricted to laser damage.

**Figure 1.**
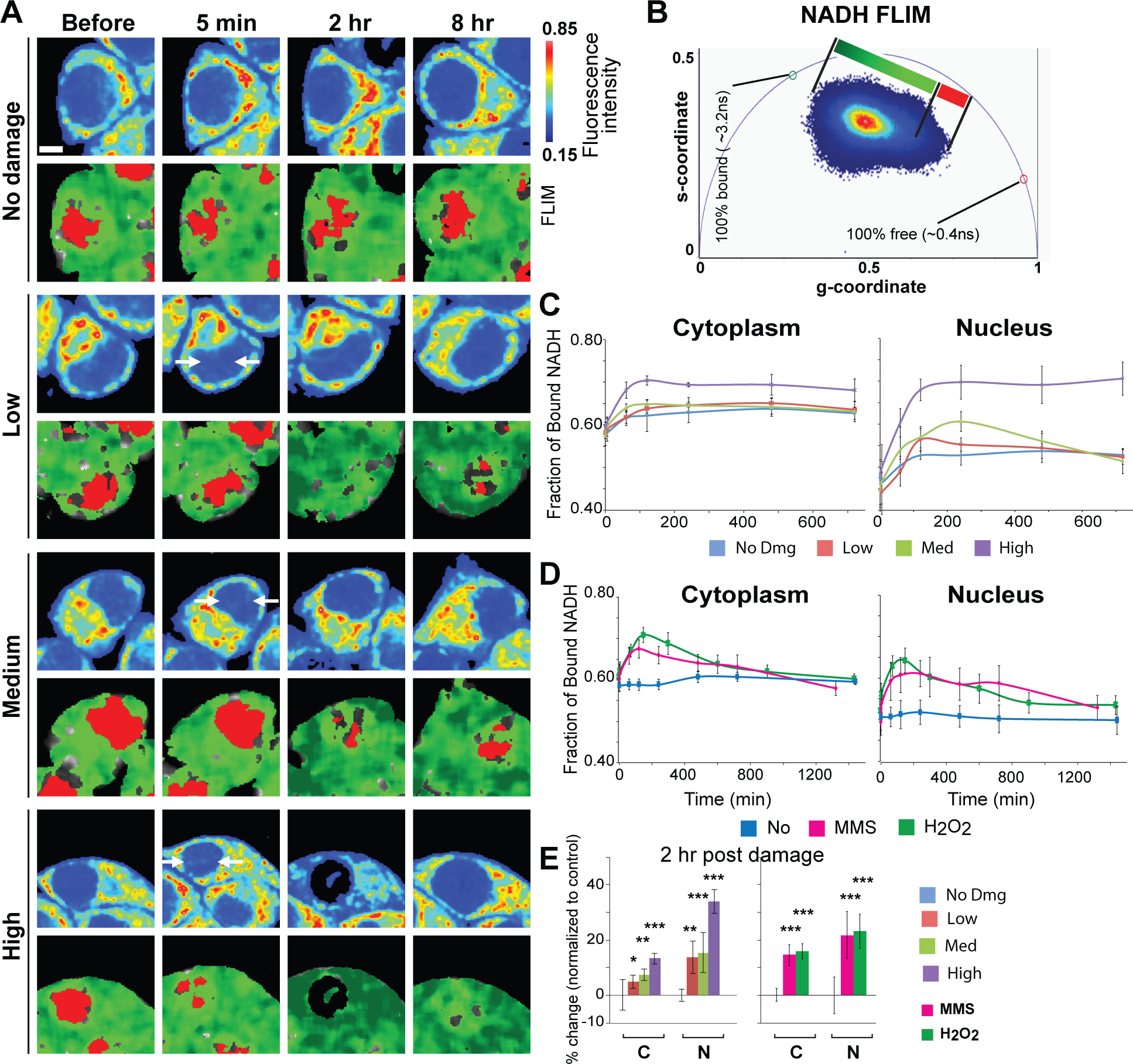
DNA damage induces rapid shift from free to bound NADH. (A) Intensity (top) and pseudo-colored FLIM (bottom) images of undamaged HeLa cells and HeLa cells damaged at low, medium, and high input laser power. In intensity images, the line color from blue to red corresponds to the normalized intensity. Damage sites are indicated by white arrows. The FLIM images are pseudo-colored according to the clusters selected on the phasor plot in (B). Scale bar = 5 µm. (B) Phasor histogram of fluorescence lifetimes on the phasor plot. Each pixel from the intensity image is transformed and plotted on the phasor diagram. Pixels are pseudo-colored based on the position along the line generated by the linear combination of short lifetime free NADH and longer lifetime protein-bound NADH. (C) The fraction of bound NADH in the cytoplasmic and nuclear compartments calculated from the position of the phasor in (B) and plotted over time. (D) The fraction of bound NADH over time in the cytoplasmic and nuclear compartments of HeLa cells treated with either 1 mM MMS or 500 µM H_2_O_2_. (E) The percent change in the fraction of bound NADH at 2 hr post damage relative to basal conditions. * p < 0.05, ** p < 0.01, *** p < 0.001.

### Increase of the bound NADH fraction is entirely dependent on PARP1 activity

What regulates the damage-induced increase of bound NADH fraction? Two major damage signaling pathways are the PARP and PIKK pathways, which are both activated in a damage dose/complexity-dependent manner (16). ATM is a member of the PIKK family, specifically activated by DNA damage to govern cell cycle checkpoint as well as DNA repair through target protein phosphorylation (41). ATM and another PIKK member DNA-PK, which mediates non-homologous end joining of DSBs, are both involved in the spreading of nuclear-wide γH2AX in response to high-dose complex DNA damage (16,42). Accordingly, PAR and γH2AX were induced initially at damage sites and spread to the whole nucleus in a damage dose-dependent manner (Fig. 2A). Thus, we examined the effect of inhibition of PARP or ATM/DNA-PK on the increase of the bound NADH fraction (Fig. 2 and Supplemental Fig. S3). As expected, inhibition of PARP by olaparib (PARPi) blocked the transient decrease of NADH intensity in both cytoplasm and nucleus, suggesting that the decrease of NADH in response to damage reflects deprivation of NAD+ by PARP, and confirmed that PARP activation originating in the nucleus affects NAD+/NADH concentration in the whole cell (Fig. 2B and C). Importantly, PARPi also dramatically suppressed the damage-induced increase of the bound NADH fraction (Fig. 2B and C). Similar results were obtained with another PARP inhibitor NU1025 (data not shown). Furthermore, comparable results were obtained with depletion of PARP1 by siRNA, indicating that PARP1, among the multiple PARP family members, is primarily responsible for the observed NADH dynamics (Fig. 2D). In contrast, ATM and DNA-PK inhibitors (AiDi) have no significant effects on either bound NADH fraction or NADH intensity (Fig. 2B and C). Taken together, the results indicate that the change in the NADH state is dependent on PARP1 activity and not ATM or DNA-PK signaling.

### Increase of bound NADH in the cytoplasm and nucleus is sensitive to mitochondrial respiratory chain complex inhibitors and rescued by NAM

The increase in protein-bound NADH in the cell was thought to reflect the oxphos pathway of energy metabolism whereas the increase in free NADH reflects the glycolytic pathway (43). Since we observed an increase of protein-bound NADH in response to damage (Fig. 1), we used the mitochondrial complex I and III respiratory (electron transport) chain inhibitors (rotenone and antimycin A, respectively) (R+A) that inhibit oxphos to examine the effect on the damage-induced increase of bound NADH species. We found that these inhibitors increased the overall basal NADH level (both intensity and concentration), which muted the response to DNA damage (Fig. 3A; data not shown). The inhibitor treatment further suppressed the damage-induced longterm increase of the protein-bound NADH fraction (observed up to 12 hr post damage) (Fig. 3A). Interestingly, the initial increase of the bound species at 1-2 hr post damage was not inhibited despite the fact that there was no significant change of NADH intensity (Fig. 3A). This suggests that the initial increase and subsequent maintenance of the high bound/free NADH ratio may be mediated by two different mechanisms.

**Figure 3.**
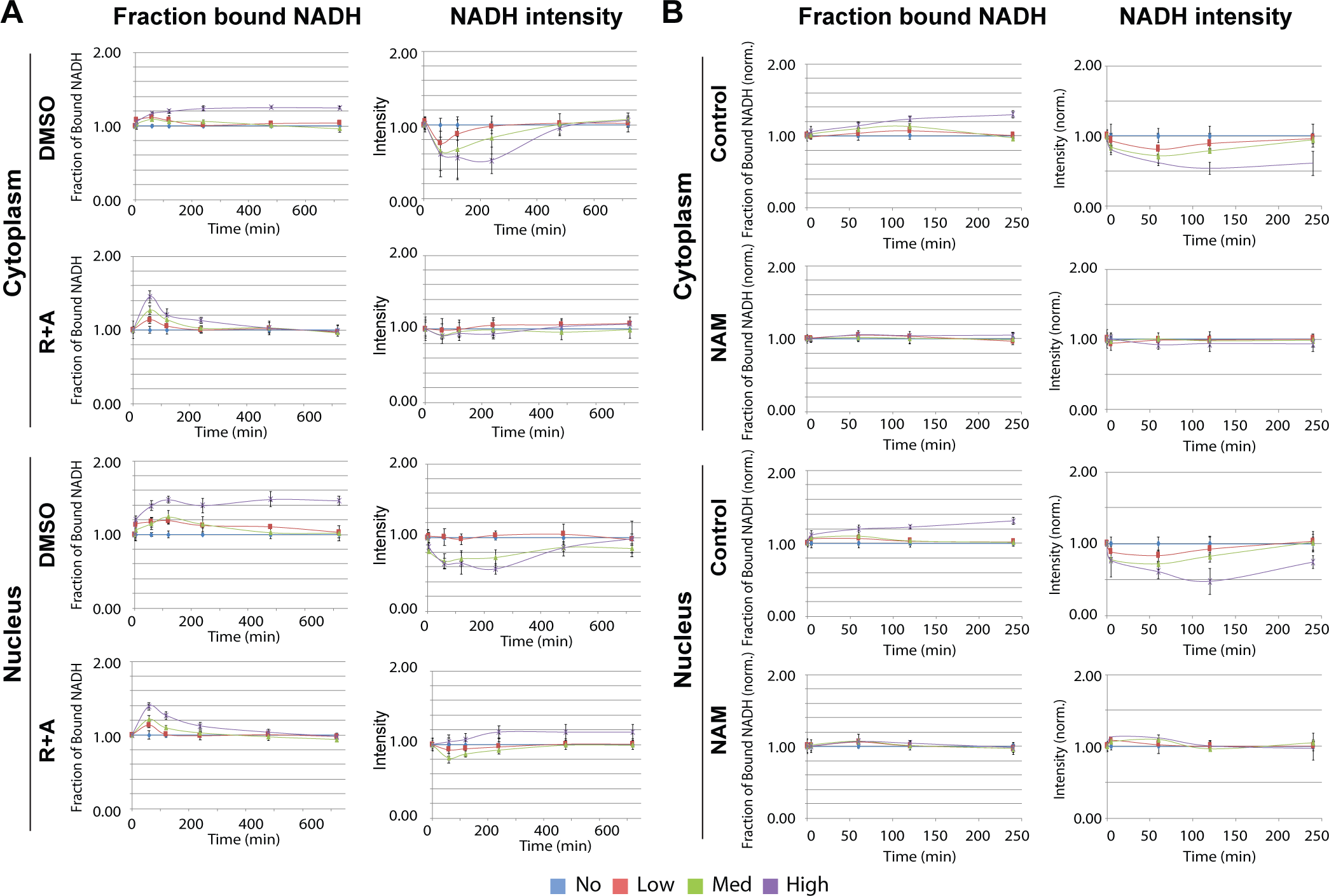
Increase of the bound NADH fraction is suppressed by the respiratory chain inhibitors. (A) The fraction of bound NADH (left) and intensity of NADH (right) in the cytoplasmic (top) and nuclear (bottom) compartments plotted over time in cells damaged with low, medium, and high input laser power in the presence of control (0.1% DMSO) or 1 µM rotenone and 1 µM antimycin A (R+A) as indicated. (B) The change in the fraction of bound NADH (left) and intensity of NADH (right) over time in cells damaged with low, medium, and high input laser power in control cells and cells pre-treated for 1 hr with 1 mM NAM.

Our analysis revealed that the increase of the bound NADH fraction occurs in both the nucleus and cytoplasm (containing mitochondria) in a highly synchronous fashion, and comparable suppression by PARP and respiratory inhibitors was observed in both subcellular regions, indicating the close communication between the two compartments (Figs. 1, 2B and 3A). The observed effect of PARP inhibition can be due to suppression of target protein PARylation and/or blocking of the intracellular NAD substrate deprivation. To test the latter hypothesis, we examined whether supplementing NAD+ would reverse the effect. Indeed, NAM, a precursor of NAD+ in the salvage pathway, not only inhibited the decrease of NADH, but also suppressed the shift to bound NADH fraction both in the nucleus and cytoplasm (Fig. 3B). The increase of bound NADH was completely suppressed during the first 4 hr unlike with R+A treatment (Fig. 3A), but comparable to PARPi (Fig. 2B). The results demonstrate that NAD+ consumption by PARP is the trigger to induce the observed shift to bound NADH.

The observed reduction of bound NADH by the respiratory chain inhibitors is concordant with the predicted inhibition of oxphos, strongly suggesting that damage-induced changes of NADH state reflect the change in energy metabolism. To further assess the relationship between the free-to-bound NADH shift and energy metabolism, metabolic flux analysis by the Seahorse XF Analyzer was performed in control and MMS-treated cells. The FLIM analyses indicate that the respiratory inhibitors have similar effects on MMS-treated cells (Figs. 3A and 4A). Namely, the partial increase of the basal NADH level and partial inhibition of the damage-induced shift from free to bound NADH were observed in both the cytoplasm and nucleus. Under this damaging condition, both oxphos as measured by the oxygen consumption rate (OCR) and glycolysis by the extracellular acidification rate (ECAR) were reduced compared to the undamaged cells, with suppression of ECAR much more severe (Fig. 4B). While PARPi has no significant effect on both basal OCR and ECAR in damaged cells, damage-induced inhibition of peak OCR and ECAR are strongly alleviated by PARPi. Cellular reliance on oxidative metabolism (OCR/ECAR) was also calculated. The results indicate the significantly elevated maximum, but not basal, oxidative reliance in MMS-treated cells compared to the control undamaged cells, which is suppressed by PARPi (Fig. 4C). Taken together, these results demonstrate that complex DNA damage that induces robust PARP activation triggers a metabolic shift resulting in greater reliance on oxphos over glycolysis.

**Figure 4.**
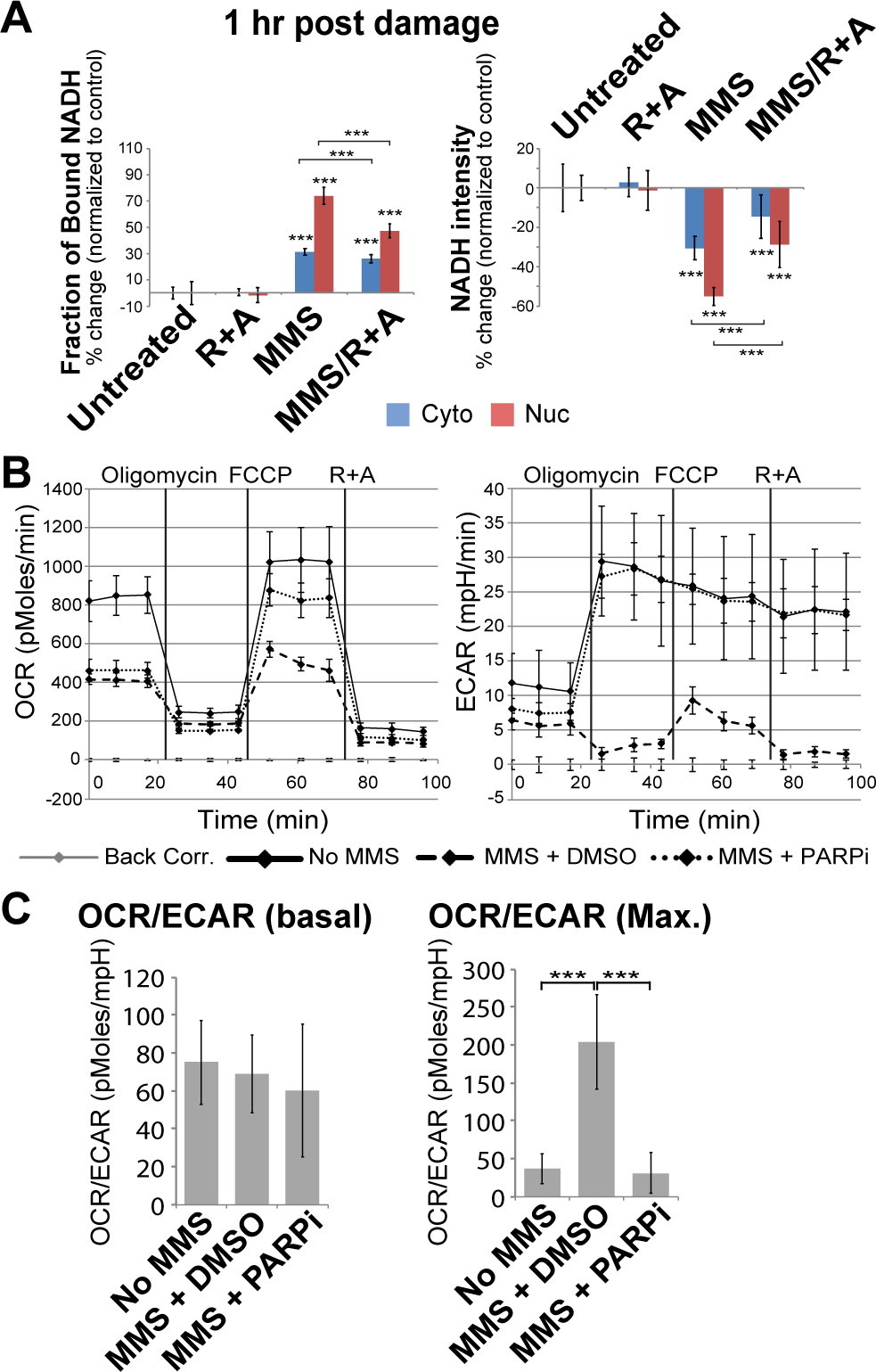
Seahorse analysis of MMS-treated cells reveal PARP-dependent increase of oxphos over glycolysis. (A) The percent change in the fraction of bound NADH (left) and intensity of NADH (right) in the cytoplasmic and nuclear compartments calculated from the position of the phasor in (Fig. 1B) and plotted over time in control undamaged cells or cells damaged with 3 mM MMS and with or without 1 µM rotenone and 1 µM antimycin A (R+A). *** p < 0.001. (B) The oxygen consumption rate (OCR, left) and extracellular acidification rate (ECAR, right) for untreated cells or cells treated with 3 mM MMS and 0.1% DMSO or 20 µM PARP inhibitor (olaparib) as analyzed by the Agilent Seahorse XF Analyzer. (C) The ratio of OCR to ECAR (oxidative reliance) evaluated at basal conditions (left) or at maximum metabolic capacities calculated following FCCP for OCR and following oligomycin for ECAR (right). *** p < 0.001.

### Oxidative phosphorylation is critical for damaged cell survival

The above results indicate that damaged cells undergo metabolic reprogramming: DNA damage increases cellular reliance on oxidative phosphorylation. Thus, we examined cellular damage sensitivity to the respiratory chain inhibitors (R+A). While the R+A treatment had no significant effect on undamaged cells, cells treated with MMS or H_2_O_2_ became significantly more sensitive to the inhibitors (Fig. 5A). This increased cytotoxicity was alleviated by PARP inhibition or PARP1 depletion, indicating that the cytotoxic effect of oxphos inhibition is PARP1-mediated (Fig. 5A, B and C). Furthermore, the increased damage sensitivity by oxphos inhibition can also be rescued by the addition of NAM (Fig. 5B). The NAM rescue effect is blocked by the inhibitor of nicotinamide phosphoribosyltransferase (NAMPT), the rate limiting enzyme for NAD+ synthesis from NAM in the salvage pathway, suggesting that the NAM conversion to NAD is required for the rescue effect (Fig. 5B and D). As expected, NAMPT inhibitor does not block the rescue effect of PARPi (Fig. 5D). Similar results were obtained with laser damage (Fig. 5E and F). Following high input-power laser damage, the survival rate of control cells at 8 hr after damage induction is approximately 83% in control cells compared to 18% in cells treated with R+A (Fig. 5E and F). This increased damage sensitivity is PARP-dependent and can be suppressed by NAM (Fig. 5E and F). Damage sensitivity was also suppressed by adding NAM even at 1 hr post laser damage induction (data not shown). Staining for PAR revealed that the amount of NAM used (1 mM) is not sufficient to inhibit PARylation, further confirming that the rescue effect of NAM is not due to PARP inhibition (Supplemental Fig. S4A). Cell death caused by respiratory inhibition of damaged cells was found to be apoptosis, and not necrosis, but is distinct from parthanatos as no AIF relocalization to the nucleus was observed (Supplemental Fig. S4B and C). Taken together, the results indicate that the oxphos pathway becomes critical for cell survival in damaged cells specifically to antagonize NAD+ depletion by PARP activation.

**Figure 5.**
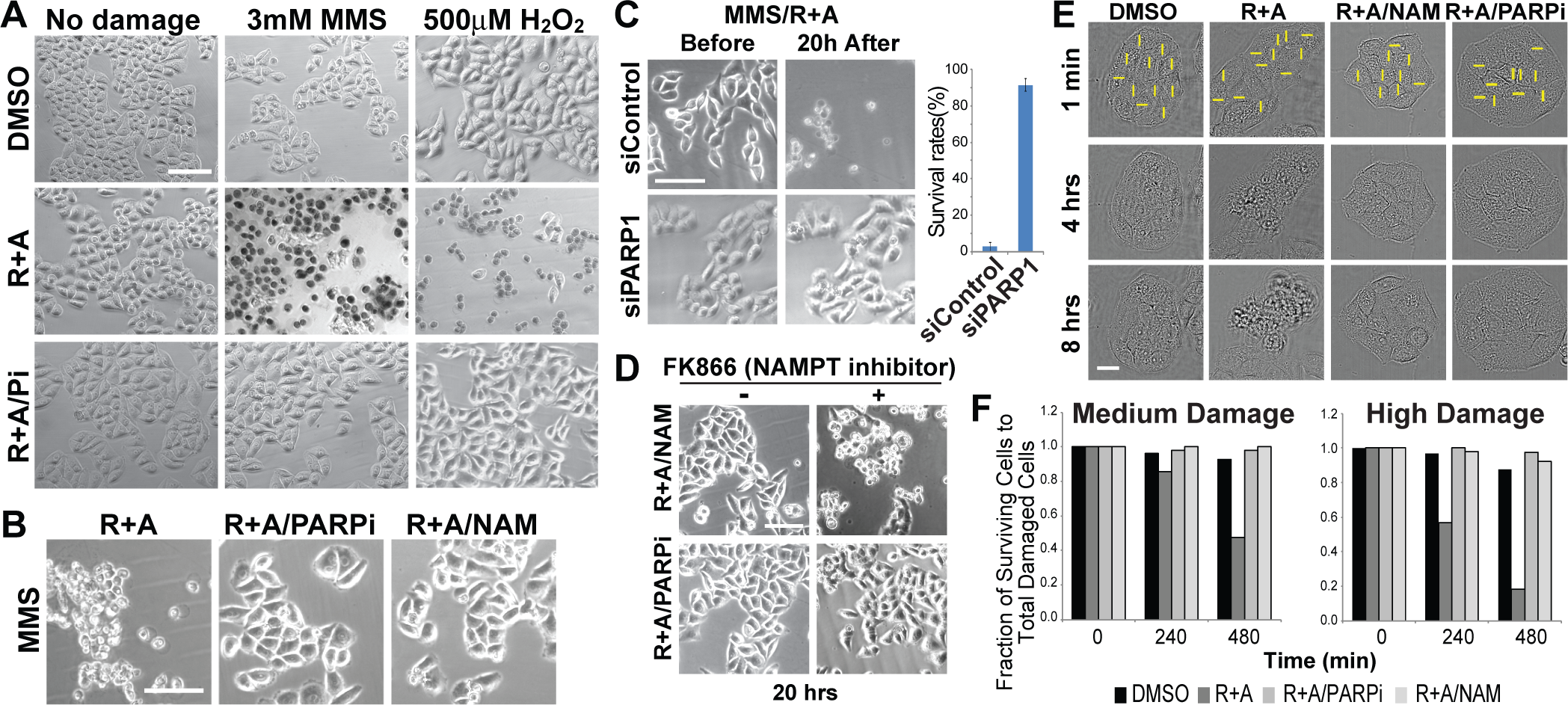
Oxphos inhibition results in increased DNA damage sensitivity, which is alleviated by PARP inhibition or NAM. (A) Cells were either undamaged or treated with MMS or H_2_O_2_ and released from damage for 20 hr in the presence of DMSO only, R+A, or R+A with PARP inhibitor (Pi). In DMSO-treated cells, no significant cytotoxicity is observed while R+A treatment resulted in significant cell death (as detected by trypan blue) only in MMS- and H_2_O_2_-treated cells. This cytotoxicity is suppressed by PARP inhibitor treatment. Scale bar = 100 µm. (B) Cytotoxic effect of R+A on MMS-treated cells (left) as in (A) is alleviated by PARPi (middle) or NAM (right). Scale bar = 100 µm. (C) The effect of control siRNA or PARP1-specific siRNA treatment in cells treated with MMS and R+A. Depletion of PARP1 (as shown in western blot in Fig. 2D) was sufficient to promote cell survival in MMS/R+A-treated cells. N=200. Scale bar = 100 µm. (D) NAM effect is NAMPT-dependent. In the presence of NAMPT inhibitor, addition of NAM failed to suppress the cytotoxicity in MMS/R+A-treated cells. PARPi-treated cells are shown for comparison. Scale bar = 100 µm. (E) Cytotoxic effect of R+A treatment in laser damage cells was suppressed by NAM or PARPi treatment. Scale bar = 20 µm. (F) Cell survival at 4 and 8 hr after laser damage induction in cells treated with DMSO, R+A, R+A with PARPi, R+A with NAM. N>50.

The observed metabolic shift in response to DNA damage is not a HeLa cell-specific phenomenon (Fig. 6). Interestingly, however, while transient NADH decrease follows similar kinetics, the extent and duration of the metabolic shift is significantly different in different cell types despite the same damage conditions (Fig. 6A, B and C). No significant shift was observed in the cytoplasm of human foreskin fibroblasts (HFF-1) cells while a prominent and persistent shift was observed in the nucleus, especially with high input-power damage (Fig. 6A). The shift in the HCT116 colorectal carcinoma cell line, in contrast, was transient (restored in less than 8 hr) and no persistent switch was observed in the nucleus even with a high input-power damage unlike HeLa or HFF-1 cells (Fig. 6B and C). HFF-1 cells are primary cells with high oxphos in contrast to highly glycolytic HCT116 cells (44,45). Furthermore, we observed that HFF-1 cells under our growth condition have significantly high basal NADH level (the basal NADH concentration is approximately five times higher in HFF-1 cells than in HCT116 cells in both the cytoplasm and nucleus) (Fig. 6A). Thus, the observed differences may reflect the differences in basal metabolic states and basal NADH concentration possibly influencing damage-induced metabolic response. Consequently, these cells exhibited different damage-induced sensitivity to respiratory inhibition (Fig. 6D). Following high input power damage, the survival rate of HCT116 cells treated with the inhibitors was approximately 51% in contrast to 87% without treatment. HFF-1 cells failed to show any increase in sensitivity to respiratory inhibition. It is possible that the amount of inhibitor used might not have been sufficient when the cells are already in the high basal oxphos state. The results reveal differential damage-induced sensitivity to oxphos inhibition in different cells, which may be dictated by the differences of basal metabolic states.

**Figure 6.**
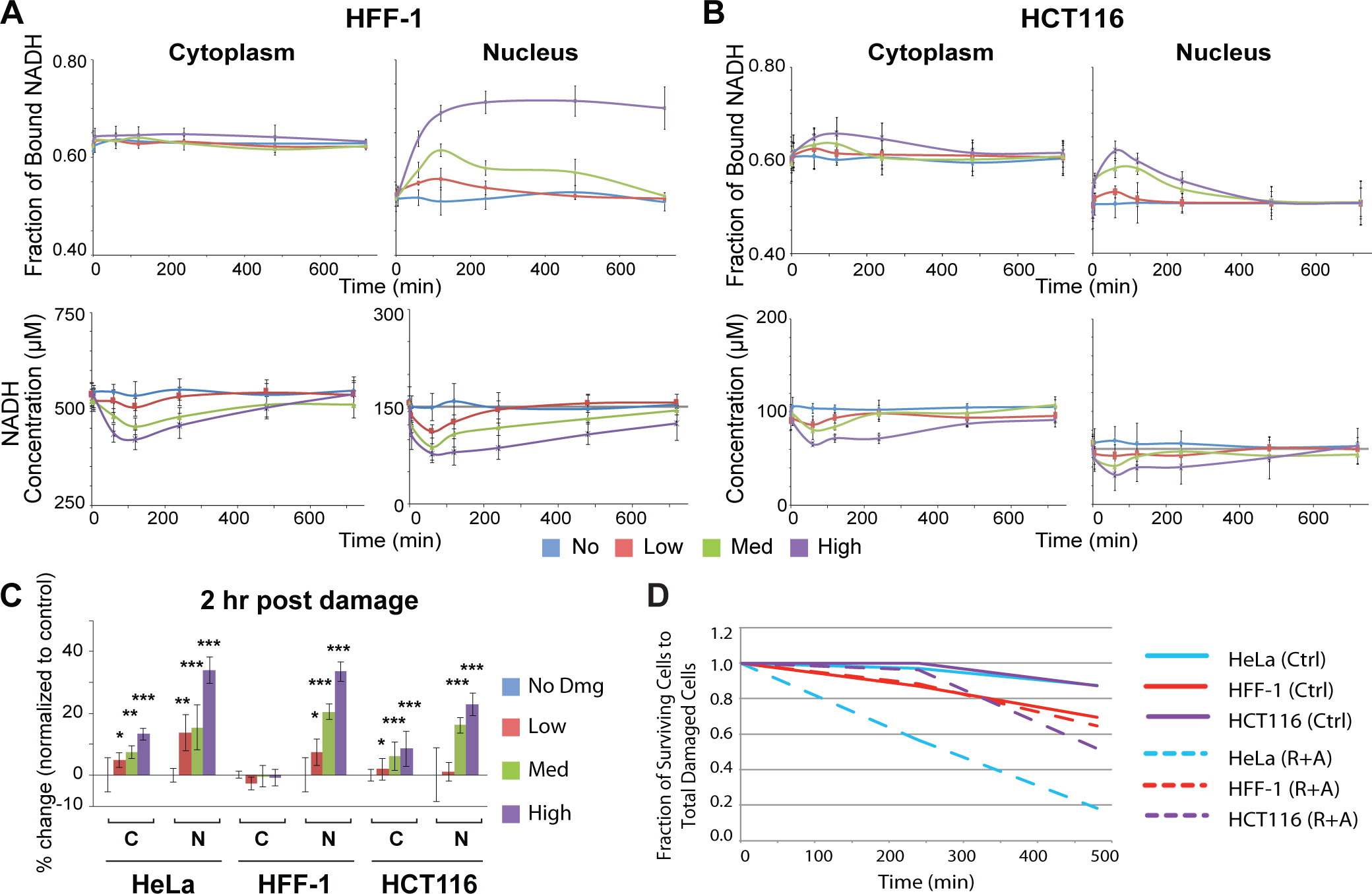
Differential free to bound NADH shift in different cell types. (A) The fraction of bound NADH in the cytoplasmic and nuclear compartments in primary HFF-1 cells following laser damage induction. Corresponding NADH concentration measurement is shown underneath. (B) The same experiments in HCT116 cells. (C) The percent change in the fraction of bound NADH in HeLa, HFF-1, and HCT116 cells at 2 hr post damage relative to basal conditions. * p < 0.05, ** p < 0.01, *** p < 0.001. (D) Differential sensitivity of HeLa, HFF-1 and HCT116 to oxphos inhibition in damaged cells. N>50.

### NAM rescues damage-induced PARP-dependent ATP depletion, but not intracellular acidification, in oxphos-inhibited cells

The above results demonstrated that damaged cells are sensitized to oxphos inhibition and either PARP1 inhibition or addition of NAM (and NAD+ generated through the salvage pathway) can rescue them. To address the mechanism, we determined ATP dynamics. DNA damage signaling and repair processes were expected to increase ATP consumption, thus reducing the intracellular ATP level. ATP would also be consumed by PARP during the PARylation reaction and was thought to be depleted as a result of PARP-mediated inhibition of glycolysis through HK leading to cell death (18,46). Using the FRET-based ATP sensors that are specifically targeted to the cytoplasm and nucleus (34), we observed the rapid reduction of ATP in the first 20 min, which persists over 6 hr post laser damage induction (Fig. 7A). ATM and DNA-PK inhibitors had a subtle effect on the initial ATP decrease within the first 20 min in both the nucleus and cytoplasm and partially reduced ATP consumption up to 6 hr in the nucleus, but not in the cytoplasm (Fig. 7A). This nuclear effect may reflect ATP consumption by DNA damage signaling and repair. In contrast, PARP inhibition completely suppressed ATP consumption in the cytoplasm and had a partial but major effect in the nucleus. The results indicate that ATP concentration in the whole cell is affected by the damage inflicted in the nucleus, and that the cytoplasmic ATP level is dictated by PARP activity.

**Figure 7.**
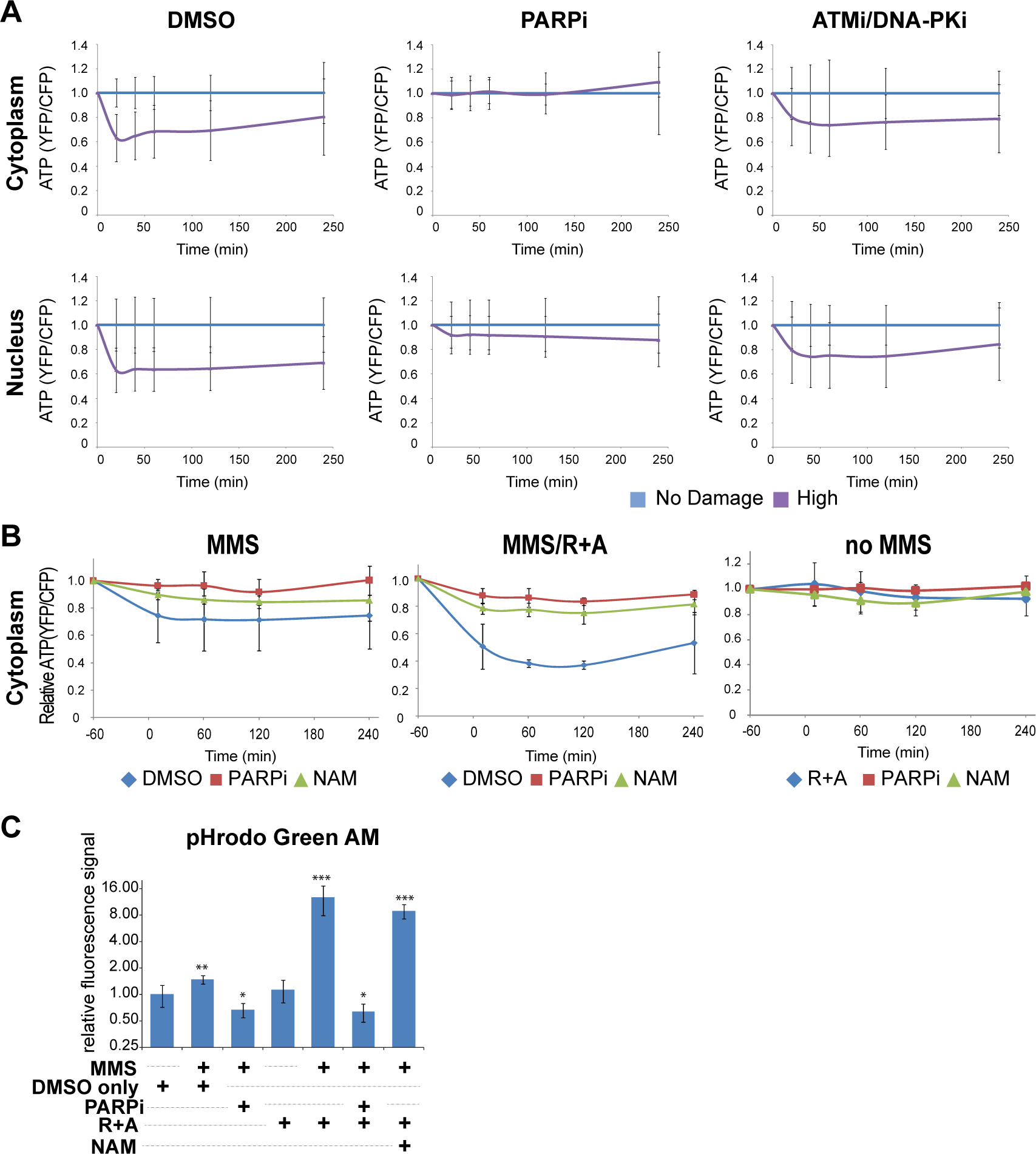
Analysis of cellular ATP and intracellular pH dynamics following DNA damage induction using ATP and pH biosensors. (A) Monitoring of relative ATP levels of HeLa cells expressing cytoplasmic (top) or nuclear (bottom) localized ATeam FRET biosensor. Cells were treated with either 0.1% DMSO (left), 20 µM PARP inhibitor (olaparib) (middle), or 10 µM ATM inhibitor (KU55933) and 10 µM DNA-PK inhibitor (NU7026) (right). Average YFP/CFP intensity was normalized with undamaged control cells at each time point. N>30. (B) Monitoring of relative cytoplasmic ATP levels of HeLa cells. Cells expressing cytoplasmic ATP sensor were cultured with medium containing with DMSO, PARPi or NAM at 1 hr before damage induction. 3 mM MMS was added at time −60 min and removed at 0 min. Averaged YFP/CFP ratio of cells was normalized with undamaged DMSO control cells at each time point. N>10. (C) HeLa cells were treated with indicated chemicals with or without MMS damage induction for 1 hr. After 2 hr, pHrodo Green AM was then added for 30 min and fluorescence intensities were measured by confocal microscopy. Data are shown as relative fluorescence intensities where higher intensities correspond to lower pH. N>25. The pHi changes were compared to cells treated with DMSO only. * p < 0.01, ** p < 0.001, *** p < 0.000001.

We next tested the effect of oxphos inhibition on the cytoplasmic ATP level in MMS-treated cells, and how PARP inhibition or addition of NAM modulates this to see if ATP deprivation can explain the oxphos inhibition-induced cytotoxicity (Fig. 7B). In undamaged cells, neither respiratory inhibition (R+A), PARPi, nor addition of NAM had any effect on the ATP level (Fig. 7B, right “no MMS”). Cytoplasmic ATP is decreased in MMS-treated cells, which is reversed by PARPi or NAM (Fig. 7B, left “MMS”). Notably, the combination of respiratory inhibition and MMS treatment resulted in significant decrease of ATP, which was also effectively alleviated by PARPi and NAM (Fig. 7B, middle “MMS/R+A”). The results reveal the damage-specific role of oxphos in ATP replenishment and confirmed that PARP activation is the major cause of cytoplasmic ATP reduction in damaged cells (Fig. 7B). The fact that addition of NAM was sufficient to also restore the ATP level strongly suggest that NAD+ is central to the maintenance of intracellular ATP in damaged cells (Fig. 7B). This is in contrast to the previous notion that PAR-dependent HK inhibition in the glycolytic pathway would result in ATP deprivation and cell death (18–20).

PARP was also shown to promote intracellular acidification, which was thought to promote cell death (23). This may possibly be due to proton production during the PARylation reaction. Thus, we also monitored intracellular pH (pHi) using the pHrodo indicator (see Methods). Although we observed a slight, but significant change in pH by MMS treatment only, we found that oxphos inhibition led to severe acidification (Fig. 7C; Supplemental Fig. S5). As expected, PARP inhibition reversed this phenomenon. Interestingly, however, NAM treatment had no significant effect on pHi, indicating that the NAM rescue effect on damage-induced sensitivity to oxphos inhibition is not due to the correction of acidic pHi, separating the two PARP-induced effects (Fig. 7C; Supplemental Fig. S5).

## Discussion

In the current study, we establish that phasor-FLIM can be used effectively to investigate real time NADH dynamics in response to DNA damage with high spatiotemporal resolution in different subcellular compartments. Our results strongly support that the increase of the protein-bound NADH fraction in response to damage can serve as an indicator for the metabolic shift to oxphos. When combined with laser microirradiation that can focus precisely inside the nucleus, the metabolic effect of nuclear damage can be specifically analyzed as opposed to the conventional damaging methods (i.e. treatment with chemical agents or whole cell irradiation) that may damage, and therefore affect function of, mitochondria. Furthermore, by regulating the laser input power, it is possible to examine the damage complexity/dosage effects on cellular metabolism. Together with using ATP and pH sensors, our results demonstrate complex metabolic consequences of damage-induced PARP1 activation in real time and uncover the crucial role of oxphos as a pro-survival effector of PARP1 signaling.

### NADH intensity/concentration and NAD+

The ratio of the photons emitted by the free and protein-bound states of NADH to total photons absorbed, known as the quantum yield, depends on the binding substrate. While the free form of NADH has a relatively low quantum yield, the protein-bound forms of NADH have a much greater quantum yield due to a decreased probability of non-radiative decay of an excited molecule while the radiative emitting pathway is largely unaffected (47,48). Thus, it is possible that typical calibrating procedures comparing intensities of known concentrations of the fluorophore to those measured in the cell may overestimate the bound NADH species. Thus, it is important to calculate and compare the absolute concentration of NADH species. We confirmed that in response to DNA damage, the changes of intensity and concentration of NADH follow similar kinetics. The concentration measurement clearly indicated comparable profiles of transient depletion of NADH despite the varying levels of basal NADH in different cell types. Importantly, our real time in situ analysis clearly indicates that although PARP1 is primarily activated in the nucleus in response to nuclear DNA damage, subsequent NADH depletion occurs in both cytoplasm and nucleus, further revealing the cell-wide metabolic effect of nucleus-initiated PARP1 activation.

Intracellular concentration of NADH was previously used to assess the single-strand break (SSB) repair capacity (as a surrogate indicator for NAD+ consumption by PARP) using a colorimetric assay of the media of a cell population (29,49). However, no direct measurement at a single cell level in response to damage has been done. In our study, NADH depletion was entirely dependent on PARP1 activity, indicating that it is a consequence of PARP1 activation in response to DNA damage. The addition of NAM, the NAD+ precursor in the salvage pathway, abolished this change without suppressing PARP activation, demonstrating that NADH reduction is due to NAD+ consumption by PARP1, rather than its PARylation activity. Interestingly, however, respiratory chain inhibition also blocked the NADH reduction despite intact PARP activation, indicating that NADH reduction in damaged cells is dependent on the mitochondrial complex activity, and may not always serve as a marker for NAD+ consumption. We attempted to measure NAD+ directly in these cells using the recently developed NAD+ sensor (32) (Supplemental Fig. S2C). However, we were unable to monitor NAD+ using this sensor in the respiratory chain inhibitor-treated cells because the sensor activity is sensitive to acidic pHi (Fig. 7C; data not shown). Nevertheless, it is clear that PARP is active (and thus, NAD+ is consumed) in these cells as additional PARP inhibition rescued the pH and cell death phenotypes. Taken together, the results strongly suggest that NAD+ depletion by activated PARP1 triggers the increased NADH consumption by the electron transport chain pathway in mitochondria (consistent with the observed increase of oxphos).

### Transient and persistent increase of bound NADH by differential PARP activation

PARP signaling was thought to be rapid and transient, affecting immediate and early response at damage sites (5,50). Indeed the PARP1 protein initially localizes at damage sites but disappears within 2 hr post damage induction (51). In addition to the localized response at damage sites, however, our results demonstrate that PARP signaling alters both NADH concentration and the free to bound NADH ratio in both the nucleus and cytoplasm (Fig. 8). NADH depletion is transient even with the high input-power damage induction, and its duration and extent are damage dose-dependent. The shift from free to bound NADH fraction is also transient in a damage dosage-dependent fashion for lower input-power damage. However, the shift becomes stabilized at high input-power damage that induces complex damage and robust PARP activation (16). This is not associated with cell death, as the majority of cells remain viable with no sign of apoptosis or necrosis during the duration of our study. Since this shift is entirely PARP-dependent, our results demonstrate that there is a threshold PARP activation, above which the metabolic change becomes persistent.

**Figure 8.**
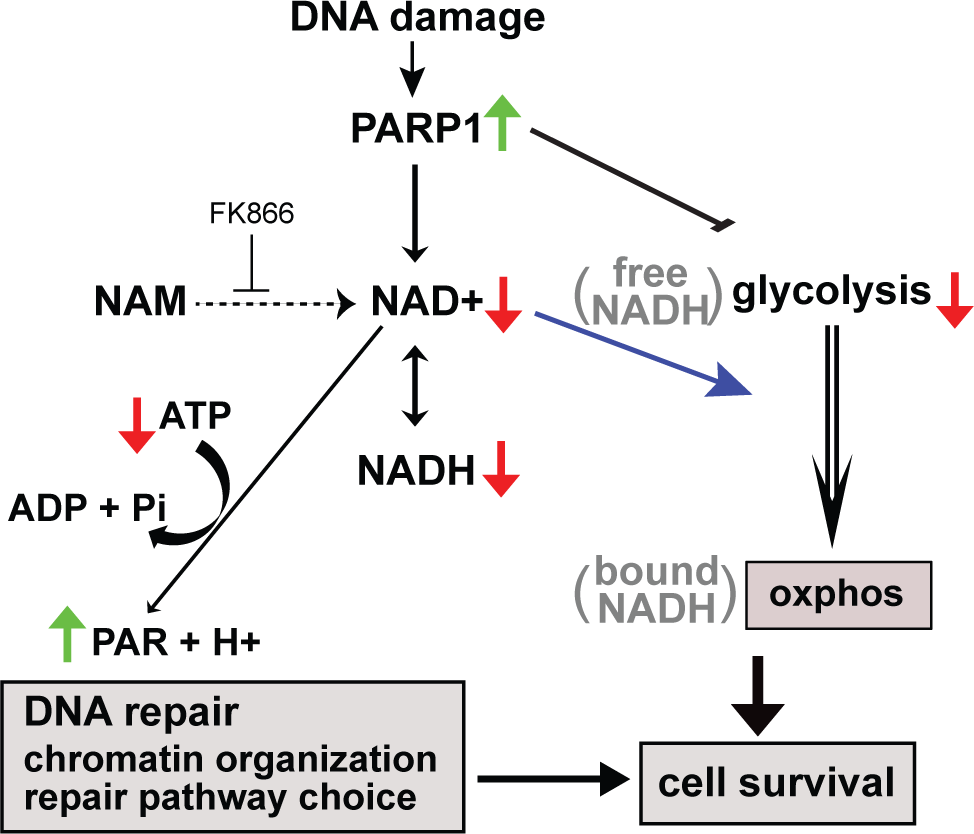
Consequences of PARP1 activation critical for damaged cell survival. DNA damage complexity/dose-dependent PARP1 activation results in accumulation of PAR at damage sites that dictate recruitment of chromatin modifiers and DNA repair pathway choice. PARylation also suppresses glycolysis through inhibition of hexokinase. NAD+ consumption by PARylation also transiently decreases NADH, which can be measured by phasor-FLIM. Decreased NAH+ also triggers the shift NADH from free to bound state, which reflects the metabolic shift from glycolysis to oxphos, respectively. Both PARP-dependent modulation of DNA repair and metabolic switch promotes damaged cell survival.

### PARP-dependent NAD+ depletion results in the shift of metabolism to oxphos

Mitochondrial respiratory chain complex inhibitors inhibit the prolonged, but not the initial, increase of bound NADH fraction, revealing the biphasic response of NADH shift to the bound form in the cell, and strongly suggesting that the persistent increase reflects a metabolic shift or reprogramming to oxphos. Seahorse experiments suggest the strong suppression of glycolysis by damage-activated PARP as reported previously (18–20). Although oxphos is also decreased by DNA damage, the extent of glycolysis suppression is greater so that there appears to be a net increase of oxphos. Consistent with this, damaged cells explicitly become sensitive to respiratory inhibition, which can be rescued by PARP1 inhibition. PARP was shown to inhibit glycolysis through PAR-mediated inhibition of the critical enzyme hexokinases resulting in energy depletion and AIF-dependent parthanatos (18). In the current study, however, the addition of NAM suppressed the shift to oxphos and rescued the cytotoxic effect of oxphos inhibition, suggesting that NAD+ depletion, and not PAR, is the critical determinant of this metabolic change critical for cell survival.

If oxphos becomes critical in damaged cells, one may predict that cells with differential metabolic states may exhibit different sensitivities to respiration inhibition. Indeed, high basal level oxphos in primary human foreskin fibroblasts (HFF) appeared to mask the change in the cytoplasm. Importantly, these cells were insensitive to the dose of respiratory inhibitors that effectively killed HeLa or HCT116 cells. Thus, our results raise an important possibility that basal respiratory activity critically determines the fate of the damaged cells with robust PARP1 activation.

## Conclusion

In summary, our results demonstrate that the phasor-FLIM method allows the real time visualization of dynamic single cell metabolic changes in response to DNA damage. With combinatorial use of laser microirradiation and fluorescence biosensors, the method was highly instrumental in uncovering the previously unrecognized long-term metabolic effect of NAD+ depletion by PARP1 activation and damage-specific role of oxphos to promote ATP production and damaged cell survival.

## Acknowledgments

This work was supported by grants from NIH P41-GM103540 (M.A.D.), NSF MCB-1615701 (M.A.D. and K.Y.), UCI Academic Senate Council on Research, Computing & Library (CORCL) (K.Y. and M.A.D.), Cancer Research Coordinating Committee (CRCC) (K.Y.), Air Force (AFOSR Grant # FA9550-08-1-0384) (M.W.B.), the Hoag Family Foundation, Huntington Beach (M.W.B.), the David and Lucille Packard Foundation, Los Altos, CA (M.W.B.), NHLBI R01 HL096987 (P.W.), and Fatima Foundation (P.W.).

The HeLa cell lines stably expressing the compartmentalized NAD+ biosensors were kindly provided by Dr. Xiaolu Cambronne at the Vollum Institute at the Oregon Health and Science University. ATP sensors were kindly provided by Dr. Hiromi Imamura at Kyoto University, Japan.

---

**Supplemental Figure S1.**
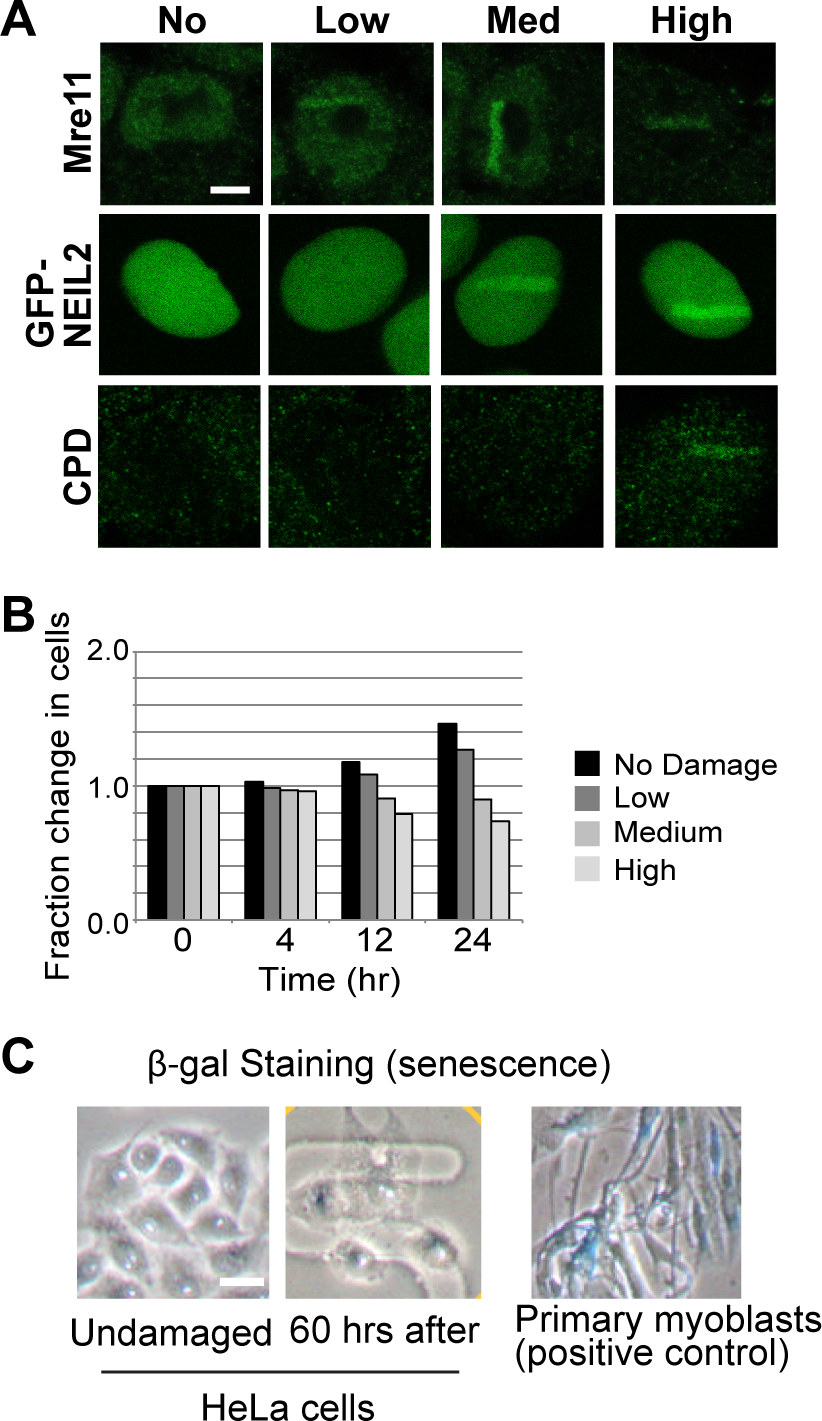
Characterization of DNA damage and cell fate following laser microirradiation using different input power. **(A)** Immuno-fluorescent staining for Mre11 (top) and CPD (bottom) in HeLa cells fixed at 1 hr post damage following low, medium, and high input laser power. Fluorescent images for HeLa cells expressing GFP-Neil2 (middle) at approximately 2 min following low, medium, and high input laser power. Scale bar = 5 µm. **(B)** The fraction change in the total number of HeLa cells over time following low, medium, and high input laser power. **(C)** (β-galactosidase staining of undamaged HeLa cells and cells at 60 hr post laser microirradiation at high input-power. Primary myoblasts were stained as a positive control. Scale bar = 20 µm.

**Supplemental Figure S2.**
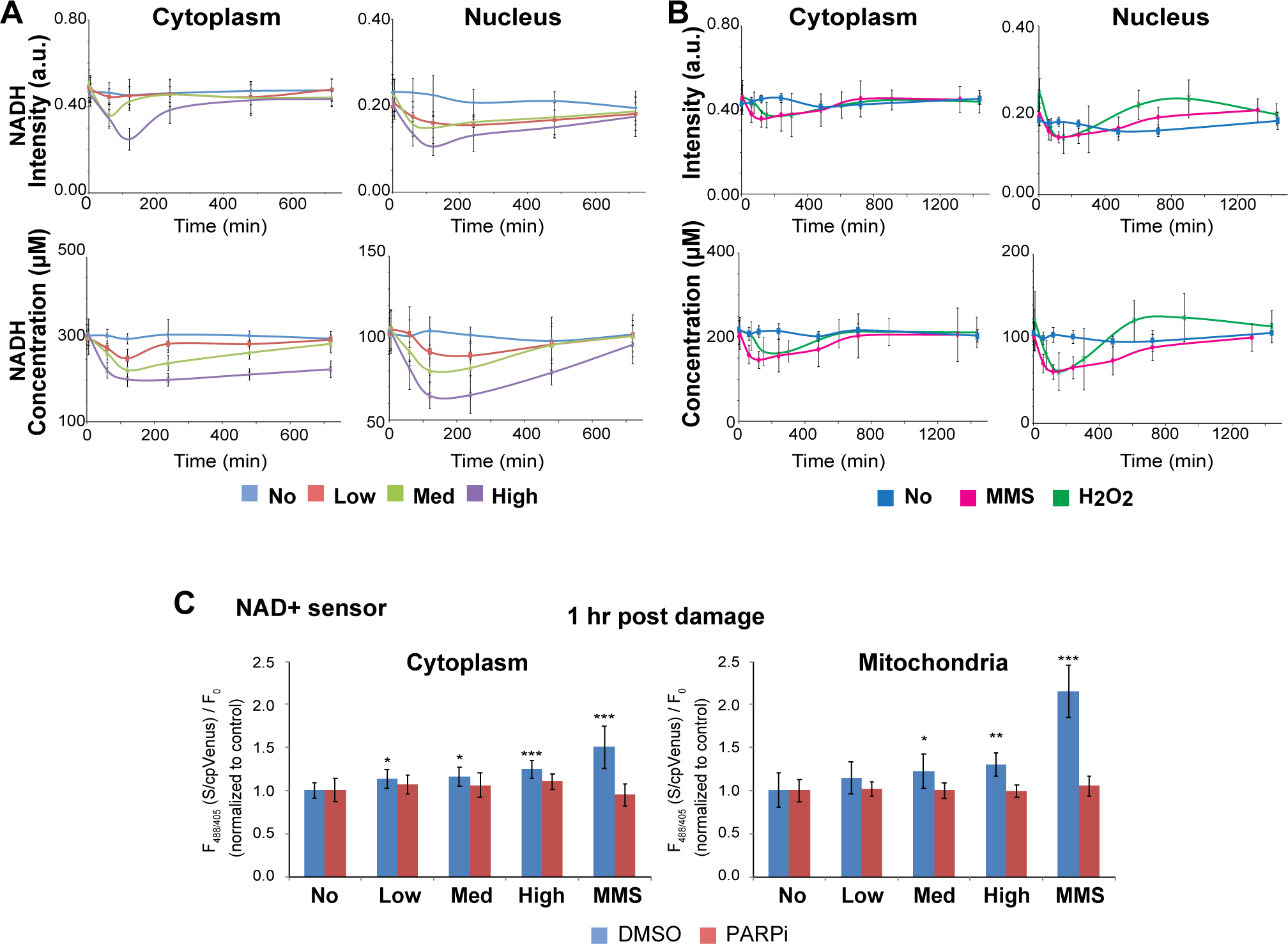
Laser Damage Induces Dose-Dependent Decrease in NADH Intensity and Concentration. **(A)** The intensity of NADH and concentration of NADH over time in the cytoplasmic and nuclear compartments of HeLa cells following low, medium, and high input laser power. **(B)** The intensity of NADH and concentration of NADH over time in the cytoplasmic and nuclear compartments of HeLa cells treated with either 1 mM MMS or 500 µM H2O2. **(C)** The change in the ratiometric fluorescence intensity of the cytoplasmic or mitochondrial NAD+ biosensor at 1 hr post damage following low, medium, and high laser microirradiation or 3 mM MMS treatment for 1 hr. Cells were treated with either 0.1% DMSO or 20 µM PARP inhibitor (olaparib). An increased fluorescence ratio reflects decreased NAD+ binding. * p < 0.05, ** p < 0.01, *** p < 0.001.

**Supplemental Figure S3.**
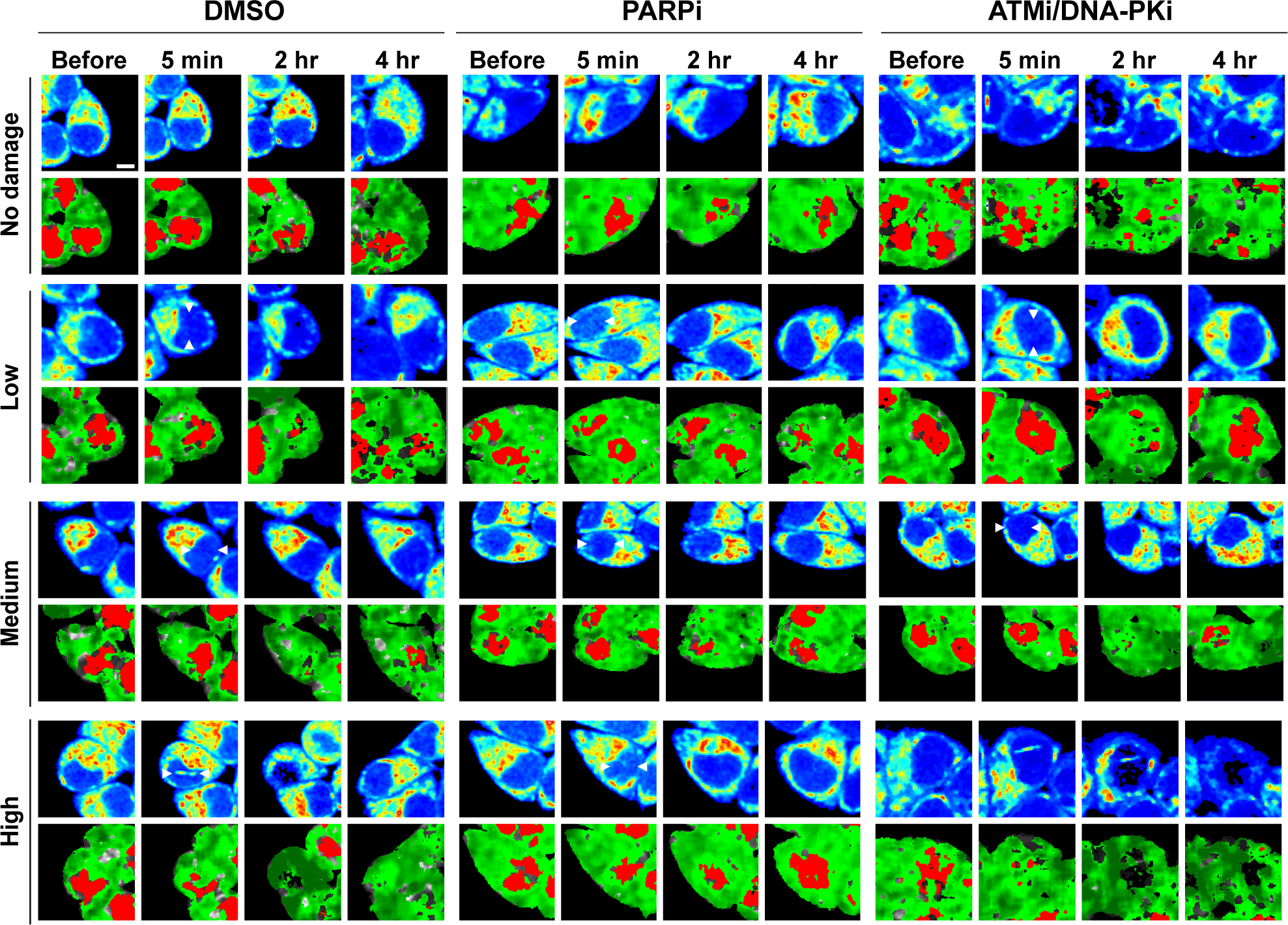
Damage-induced changes in NADH intensity and FLIM images are PARP, but not ATM/DNA-PK,-dependent. Intensity (top) and pseudo-colored FLIM (bottom) images of undamaged HeLa cells and HeLa cells damaged at low, medium, and high input laser power and treated with either 0.1% DMSO (left), 20 µM PARP inhibitor (olaparib) (middle), or 10 µM ATM inhibitor (KU55933) and 10 µM DNA-PK inhibitor (NU7026) (right). In intensity images, the line color from blue to red corresponds to the normalized intensity. Damage sites are indicated by white arrows. The FLIM images are pseudo-colored according to the clusters selected on the phasor plot in (Figure 1B). Scale bar = 5 µm.

**Supplemental Figure S4.**
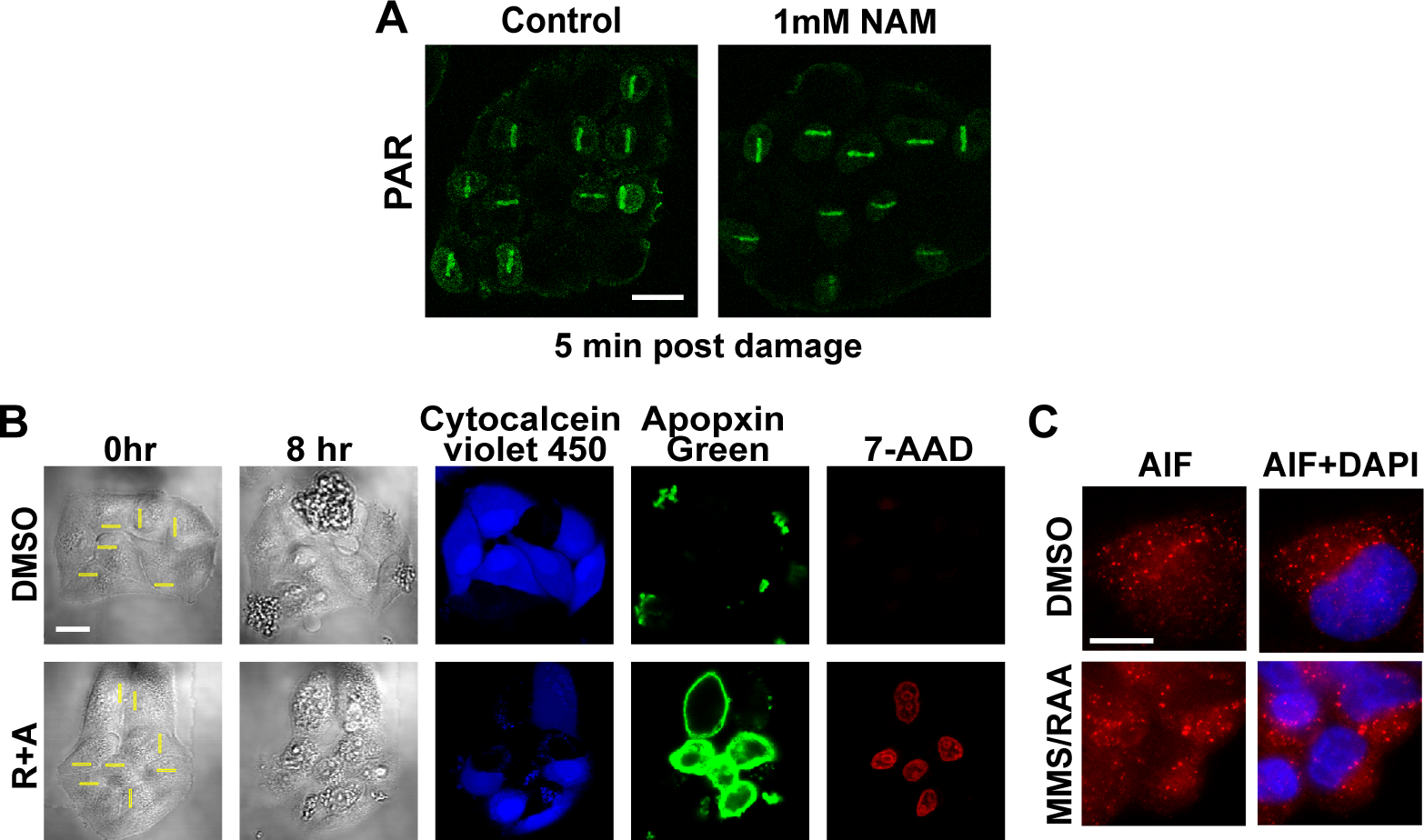
Damage-specific cell death induced by R+A treatment is AIF-independent apoptosis. **(A)** Immuno-fluorescent staining for PAR (green) in HeLa cells pre-treated with or without 1 mM NAM for 1 hr and fixed 5 min post damage following low, medium, and high input laser power. Scale bar = 20 µm. **(B)** Brightfield images of HeLa cells treated with either 0.2 % DMSO or 1 µM rotenone and 1 µM antimycin A before damage and 8 hr post damage. Apoptosis and necrosis was detected using a commercial kit (AB176749) for live cells (cytocalcein violet 450, blue), phosphatidylserine exposure (apopxin green, green), and loss of plasma integrity (7-aminoactinomycin D, red). Scale bar = 20 µm. **(C)** Immuno-fluorescent staining for apoptosis inducing factor (AIF) and DAPI in HeLa cells treated with either 0.1% DMSO or 1 mM MMS, 1 µM rotenone, and 1 µM antimycin A. Scale bar = 10 µm.

**Supplemental Figure S5.**
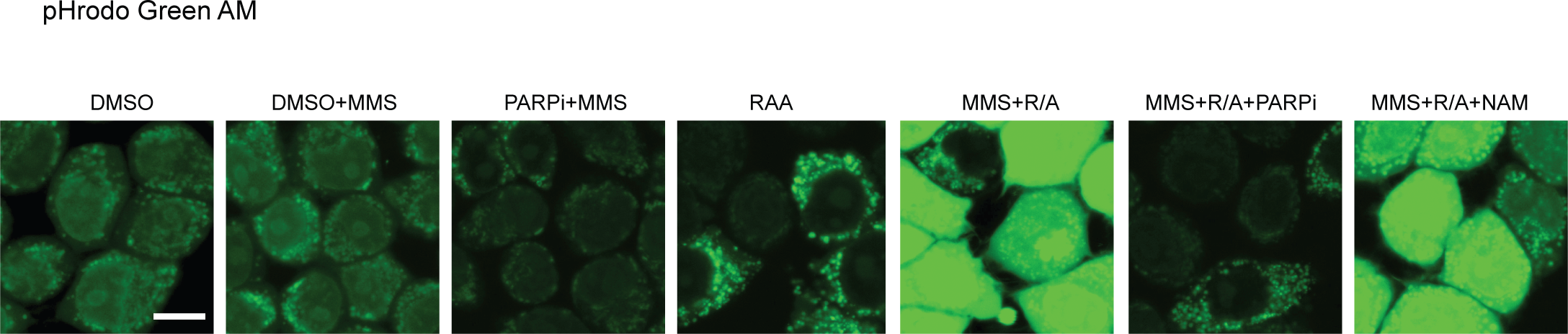
Intracellular pH (pHi) measurement using pHrodo Green AM reveal significant damage-specific acidification by R+A, which is alleviated by PARPi but not by NAM. Fluorescence images of HeLa cells were treated with indicated chemicals with or without MMS damage induction for 1 hr with pHrodo Green AM. Relative fluorescence intensities were quantified in Fig. 7C where higher intensities correspond to lower pH. Scale bar = 10 µm.

## References

1. Gupte, R., Liu, Z. and Kraus, W.L. (2017) PARPs and ADP-ribosylation: recent advances linking molecular functions to biological outcomes. Genes Dev., 31, 101–126.

2. Ahel, D., Horejsí, Z., Wiechens, N., Polo, S.E., Garcia-Wilson, E., Ahel, I., Flynn, H., Skehel, M., West, S.C., Jackson, S.P. et al. (2009) Poly(ADP-ribose)-dependent regulation of DNA repair by the chromatin remodeling enzyme ALC1. Science, 325, 1240–1243.

3. Altmeyer, M., Neelsen, K.J., Teloni, F., Pozdnyakova, I., Pellegrino, S., Grøfte, M., Rask, M.B., Streicher, W., Jungmichel, S., Nielsen, M.L. et al. (2015) Liquid demixing of intrinsically disordered proteins is seeded by poly(ADP-ribose). Nat. Commun., 6, 8088.

4. Ayrapetov, M.K., Gursoy-Yuzugullu, O., Xu, C., Xu, Y. and Price, B.D. (2014) DNA double-strand breaks promote methylation of histone H3 on lysine 9 and transient formation of repressive chromatin. Proc. Natl. Acad. Sci., 111, 9169–9174.

5. Ball, A.R., Jr. and Yokomori, K. (2011) Damage site chromatin: open or closed? Curr. Opin. Cell Biol., 23, 277–283.

6. Bouwman, P., Aly, A., Escandell, J.M., Pieterse, M., Bartkova, J., van der Gulden, H., Hiddingh, S., Thanasoula, M., Kulkarni, A., Yang, Q. et al. (2010) 53BP1 loss rescues BRCA1 deficiency and is associated with triple-negative and BRCA-mutated breast cancers. Nat. Struc. Mol. Biol., 17, 688–695.

7. Chou, D.M., Adamson, B., Dephoure, N.E., Tan, X., Nottke, A.C., Hurov, K.E., Gygi, S.P., Colaiácovo, M.P. and Elledge, S.J. (2010) A chromatin localization screen reveals poly (ADP ribose)-regulated recruitment of the repressive polycomb and NuRD complexes to sites of DNA damage. Proc. Natl. Acad. Sci., 107, 18475–18480.

8. Gottschalk, A.J., Timinszky, G., Kong, S.E., Jin, J., Cai, Y., Swanson, S.K., Washburn, M.P., Florens, L., Ladurner, A.G., Conaway, J.W. et al. (2009) Poly(ADP-ribosyl)ation directs recruitment and activation of an ATP-dependent chromatin remodeler. Proc. Natl. Acad. Sci., 106, 13770–13774.

9. Izhar, L., Adamson, B., Ciccia, A., Lewis, J., Pontano-Vaites, L., Leng, Y., Liang, A.C., Westbrook, T.F., Harper, J.W. and Elledge, S.J. (2015) A Systematic Analysis of Factors Localized to Damaged Chromatin Reveals PARP-Dependent Recruitment of Transcription Factors. Cell Rep., 11, 1486–1500.

10. Jaspers, J.E., Kersbergen, A., Boon, U., Sol, W., van Deemter, L., Zander, S.A., Drost, R., Wientjens, E., Ji, J., Aly, A. et al. (2013) Loss of 53BP1 causes PARP inhibitor resistance in Brca1-mutated mouse mammary tumors. Cancer Discov., 3, 68–81.

11. Khoury-Haddad, H., Guttmann-Raviv, N., Ipenberg, I., Huggins, D., Jeyasekharan, A.D. and Ayoub, N. (2014) PARP1-dependent recruitment of KDM4D histone demethylase to DNA damage sites promotes double-strand break repair. Proc. Natl. Acad. Sci., 111, E728–737.

12. Larsen, D.H., Poinsignon, C., Gudjonsson, T., Dinant, C., Payne, M.R., Hari, F.J., Danielsen, J.M., Menard, P., Sand, J.C., Stucki, M. et al. (2010) The chromatin-remodeling factor CHD4 coordinates signaling and repair after DNA damage. J. Cell Biol., 190, 731–740.

13. Polo, S.E., Kaidi, A., Baskcomb, L., Galanty, Y. and Jackson, S.P. (2010) Regulation of DNA-damage responses and cell-cycle progression by the chromatin remodelling factor CHD4. EMBO J., 29, 3130–3139.

14. Smeenk, G., Wiegant, W.W., Vrolijk, H., Solari, A.P., Pastink, A. and van Attikum, H. (2010) The NuRD chromatin-remodeling complex regulates signaling and repair of DNA damage. J. Cell Biol., 190, 741–749.

15. Sun, Y., Jiang, X., Xu, Y., Ayrapetov, M.K., Moreau, L.A., Whetstine, J.R. and Price, B.D. (2009) Histone H3 methylation links DNA damage detection to activation of the tumour suppressor Tip60. Nat. Cell Biol., 11, 1376–1382.

16. Cruz, G.M.S., Kong, X., Silva, B.A., Khatibzadeh, N., Thai, R., Berns, M.W. and Yokomori, K. (2015) Femtosecond near-infrared laser microirradiation reveals a crucial role for PARP signaling on factor assemblies at DNA damage sites. Nuc. Acids Res., 44, e27.

17. Heikal, A.A. (2010) Intracellular coenzymes as natural biomarkers for metabolic activities and mitochondrial anomalies. Biomark. Med., 4, 241–263.

18. Andrabi, S.A., Umanah, G.K., Chang, C., Stevens, D.A., Karuppagounder, S.S., Gagné, J.P., Poirier, G.G., Dawson, V.L. and Dawson, T.M. (2014) Poly(ADP-ribose) polymerase-dependent energy depletion occurs through inhibition of glycolysis. Proc. Natl. Acad. Sci., 111, 10209–10214.

19. Feng, F.Y., de Bono, J.S., Rubin, M.A. and Knudsen, K.E. (2015) Chromatin to Clinic: The Molecular Rationale for PARP1 Inhibitor Function. Mol. Cell, 58, 925–934.

20. Fouquerel, E., Goellner, E.M., Yu, Z., Gagne, J.P., Barbi de Moura, M., Feinstein, T., Wheeler, D., Redpath, P., Li, J., Romero, G. et al. (2014) ARTD1/PARP1 negatively regulates glycolysis by inhibiting hexokinase 1 independent of NAD+ depletion. Cell Rep, 8, 1819–1831.

21. Andrabi, S.A., Dawson, T.M. and Dawson, V.L. (2008) Mitochondrial and nuclear cross talk in cell death: parthanatos. Ann. N. Y. Acad. Sci., 1147, 233–241.

22. Fatokun, A.A., Dawson, V.L. and Dawson, T.M. (2014) Parthnatos: mitochondrial-linked mechanisms and therapeutic opportunities. Br. J. Pharmacol., 171, 2000–2016.

23. Affar, E.B., Shah, R.G., Dallaire, A.K., Castonguay, V. and Shah, G.M. (2002) Role of poly(ADP-ribose) polymerase in rapid intracellular acidification induced by alkylating DNA damage. Proc. Natl. Acad. Sci., 99, 245–250.

24. Stringari, C., Edwards, R.A., Pate, K.T., Waterman, M.L., Donovan, P.J. and Gratton, E. (2012) Metabolic trajectory of cellular differentiation in small intestine by Phasor Fluorescence Lifetime Microscopy of NADH. Sci. Rep., 2, 568.

25. Stringari, C., Sierra, R., Donovan, P.J. and Gratton, E. (2012) Label-free separation of human embryonic stem cells and their differentiating progenies by phasor fluorescence lifetime microscopy. J. Biomed. Opt., 17, 046012.

26. Skala, M. and Ramanujam, N. (2010) Multiphoton redox ratio imaging for metabolic monitoring in vivo. Methods Mol Biol, 594, 155–162.

27. Digman, M.A., Caiolfa, V.R., Zamai, M. and Gratton, E. (2008) The phasor approach to fluorescence lifetime imaging analysis. Biophys. J., 94, L14–16.

28. Ma, N., Digman, M.A., Malacrida, L. and Gratton, E. (2016) Measurements of absolute concentrations of NADH in cells using the phasor FLIM method. Biomed. Opt. Express, 7, 2441–2452.

29. Nakamura, J., Asakura, S., Hester, S.D., de Murcia, G., Caldecott, K.W. and Swenberg, J.A. (2003) Quantitation of intracellular NAD(P)H can monitor an imbalance of DNA single strand break repair in base excision repair deficient cells in real time. Nuc. Acids Res., 31, e104.

30. Heale, J.T., Ball, J., A. R., Schmiesing, J.A., Kim, J.S., Kong, X., Zhou, S., Hudson, D., Earnshaw, W.C. and Yokomori, K. (2006) Condensin I interacts with the PARP-1-XRCC1 complex and functions in DNA single-stranded break repair. Mol. Cell, 21, 837–848.

31. Wright, B.K., Andrews, L.M., Markham, J., Jones, M.R., Stringari, C., Digman, M.A. and Gratton, E. (2012) NADH distribution in live progenitor stem cells by phasor-fluorescence lifetime image microscopy. Biophys. J., 103, L7–9.

32. Cambronne, X.A., Stewart, M.L., Kim, D., Jones-Brunette, A.M., Morgan, R.K., Farrens, D.L., Cohen, M.S. and Goodman, R.H. (2016) Biosensor reveals multiple sources for mitochondrial NAD^+^. Science, 352, 1474–1477.

33. Kioka, H., Kato, H., Fujikawa, M., Tsukamoto, O., Suzuki, T., Imamura, H., Nakano, A., Higo, S., Yamazaki, S., Matsuzaki, T. et al. (2014) Evaluation of intramitochondrial ATP levels identifies G0/G1 switch gene 2 as a positive regulator of oxidative phosphorylation. Proc. Natl. Acad. Sci., 111, 273–278.

34. Imamura, H., Nhat, K.P., Togawa, H., Saito, K., Iino, R., Kato-Yamada, Y., Nagai, T. and Noji, H. (2009) Visualization of ATP levels inside single living cells with fluorescence resonance energy transfer-based genetically encoded indicators. Proc. Natl. Acad. Sci., 106, 15651–15656.

35. Chen, Y., Chernyavsky, A., Webber, R.J., Grando, S.A. and Wang, P.H. (2015) Critical role of the neonatal Fc receptor (FcRn) in the pathogenic action of antimitochondrial autoantibodies synergizing with anti-desmoglein autoantibodies in pemphigus vulgaris. J. Biol. Chem., 290, 23826–23837.

36. Gey, C. and Seeger, K. (2013) Metabolic changes during cellular senescence investigated by proton NMR-spectroscopy. Mech. Ageing Dev., 134, 130–138.

37. Gomez-Godinez, V., Wu, T., Sherman, A.J., Lee, C.S., Liaw, L.H., Zhongsheng, Y., Yokomori, K. and Berns, M.W. (2010) Analysis of DNA double-strand break response and chromatin structure in mitosis using laser microirradiation. Nuc. Acids Res., 38, e202.

38. Kong, X., Mohanty, S.K., Stephens, J., Heale, J.T., Gomez-Godinez, V., Shi, L.Z., Kim, J.S., Yokomori, K. and Berns, M.W. (2009) Comparative analysis of different laser systems to study cellular responses to DNA damage in mammalian cells. Nucleic Acids Res., 37, e68.

39. Kong, X., Stephens, J., Ball, A.R., Jr., Heale, J.T., Newkirk, D.A., Berns, M.W. and Yokomori, K. (2011) Condensin I Recruitment to Base Damage-Enriched DNA Lesions Is Modulated by PARP1. PLoS One, 6, e23548.

40. Silva, B.A., Stambaugh, J.R., Yokomori, K., Shah, J.V. and Berns, M.W. (2014) DNA damage to a single chromosome end delays anaphase onset. J. Biol. Chem., 289, 22771–22784.

41. Shiloh, Y. (2003) ATM and related protein kinases: safeguarding genome integrity. Nat. Rev. Cancer, 3, 155–168.

42. Burma, S., Chen, B.P., Murphy, M., Kurimasa, A. and Chen, D.J. (2001) ATM phosphorylates histone H2AX in response to DNA double-strand breaks. J. Biol. Chem., 276, 42462–42467.

43. Stringari, C., Cinquin, A., Cinquin, O., Digman, M.A., Donovan, P.J. and Gratton, E. (2011) Phasor approach to fluorescence lifetime microscopy distinguishes different metabolic states of germ cells in a live tissue. Proc. Natl. Acad. Sci., 108, 13582–13587.

44. Zheng, J. (2012) Energy metabolism of cancer: Glycolysis versus oxidative phosphorylation (Review). Oncol. Lett., 4, 1151–1157.

45. Nickens, K.P., Wikstrom, J.D., Shirihai, O.S., Patierno, S.R. and Ceryak, S. (2013) A bioenergetic profile of non-transformed fibroblasts uncovers a link between death-resistance and enhanced spare respiratory capacity. Mitochondrion, 13, 662–667.

46. Fouquerel, E., Goellner, E.M., Yu, Z., Gagné, J.P., Barbi de Moura, M., Feinstein, T., Wheeler, D., Redpath, P., Li, J., Romero, G. et al. (2014) ARTD1/PARP1 negatively regulates glycolysis by inhibiting hexokinase 1 independent of NAD+ depletion. Cell Rep., 8, 1819–1831.

47. Scott, T.G., Spencer, R.D., Leonard, N.J. and Weber, G. (1970) Synthetic spectroscopic models related to coenzymes and base pairs. V. Emission properties of NADH. Studies of fluorescence lifetimes and quantum efficiencies of NADH, AcPyADH, [reduced acetylpyridineadenine dinucleotide] and simplified synthetic models. J. Am. Chem. Soc., 92, 687–695.

48. Vishwasrao, H.D., Heikal, A.A., Kasischke, K.A. and Webb, W.W. (2005) Conformational dependence of intracellular NADH on metabolic state revealed by associated fluorescence anisotropy. J. Biol. Chem., 280, 25119–25126.

49. Yoshimura, M., Kohzaki, M., Nakamura, J., Asagoshi, K., Sonoda, E., Hou, E., Prasad, R., Wilson, S.H., Tano, K., Yasui, A. et al. (2006) Vertebrate POLQ and POLbeta cooperate in base excision repair of oxidative DNA damage. Mol. Cell, 24, 115–125.

50. Polo, S.E. and Jackson, S.P. (2011) Dynamics of DNA damage response proteins at DNA breaks: a focus on protein modifications. Genes Dev., 25, 409–433.

51. Kim, J.-S., Krasieva, T.B., Kurumizaka, H., Chen, D.J., Taylor, A.M. and Yokomori, K. (2005) Independent and sequential recruitment of NHEJ and HR factors to DNA damage sites in mammalian cells. J. Cell Biol., 170, 341–347.

